# Taxonomic turnover with phylogenetic conservation in subaerial biofilm (SAB) communities of marble monuments

**DOI:** 10.64898/2026.11.13.766465

**Authors:** Jean-David Grattepanche, Judith M Jacob, Alberto Tenore, Fabiana Russo, Bettina Buttaro, Robert W Sanders, Isaac Klapper

**Author notes:** Corresponding author: Jean-David Grattepanche. All authors reviewed and approved the final manuscript.

## Abstract

Subaerial biofilms (SABs) colonizing exposed stone surfaces provide natural systems for investigating microbial community assembly under challenging environmental conditions. Although bacterial diversity on stone has been widely described, whether taxonomic variation reflects broader phylogenetic and ecological differences remains unclear. Here, we characterized SAB communities inhabiting two geographically separated historic urban marble monuments, Federal Hall National Memorial (New York) and the Lincoln Memorial (Washington, DC), using SSU rRNA amplicon sequencing. Additional samples from the Merchants’ Exchange Building (Philadelphia) were also examined. Bacterial communities exhibited pronounced taxonomic turnover but substantially greater phylogenetic similarity, indicating that community variation primarily reflected replacement among closely related bacterial lineages rather than turnover among distantly related clades. Despite this taxonomic turnover, core assemblage analyses identified conserved bacterial lineages shared across monuments, several of which were also detected in the independent Philadelphia samples. Consistent with these patterns, inferred ecological composition remained remarkably similar among monuments, with chlorophyll-containing bacteria dominating all communities. Together, these findings demonstrate that these monuments harbor distinct bacterial assemblages while maintaining a conserved phylogenetic and inferred ecological framework, suggesting that environmental filtering and local ecological processes may contribute to SAB community assembly across spatial scales.

## Introduction

Biofilms are structured microbial communities that develop on surfaces, where interactions among microorganisms and their surrounding environment influence community assembly and persistence. On exposed terrestrial surfaces, subaerial biofilms (SABs) experience fluctuating environmental conditions, including variations in moisture availability, temperature, and solar radiation (Gorbushina & Broughton, 2009; Villa & Cappitelli, 2019; Tenore et al., 2024). These communities can act as pioneer colonizers of exposed mineral surfaces, sometimes preceding lichen establishment (Gorbushina, 2007). Through biological activity, SABs can alter the physical and chemical properties of rock surfaces and exhibit heterogeneous patterns of colonization (Gorbushina, 2007). Despite exposure to desiccation, intense radiation, and nutrient limitation, SABs can persist through interactions among phototrophic and heterotrophic microorganisms (Gorbushina & Broughton, 2009; Tenore et al., 2024).

The composition and physical characteristics of stone substrates, together with local environmental conditions, can influence the colonization and community structure of SABs (Villa et al., 2016). Studies of SABs on historic monuments and outdoor stone surfaces have highlighted the importance of environmental exposure and surface characteristics. In particular, differences in surface orientation, exposure to sunlight, and stone geometry have been associated with variations in microbial colonization and community structure on sandstone and limestone sculptures (Cattò et al., 2023). On marble monuments, moisture availability and temporal environmental variability may also influence the persistence and activity of SAB communities (Tenore et al., 2024). Together, these studies suggest that substrate characteristics and local environmental conditions contribute to spatial heterogeneity in SAB communities, potentially even among neighboring surfaces of the same monument.

Microbial communities exhibit biogeographic patterns shaped by the combined effects of environmental selection, dispersal, ecological drift, and evolutionary processes (Hanson et al., 2012). These patterns have been documented across diverse environments, where geographic separation and environmental heterogeneity are associated with differences in microbial community composition. On exposed stone surfaces, Brewer and Fierer (2018) reported associations between bacterial and eukaryotic communities and geographic location, climate, and substrate composition across limestone and granite tombstones sampled on three continents. More recently, Trovão et al. (2024) reported spatial differences in microbial community composition across locations on a marble statue, highlighting the importance of local spatial organization. At a broader geographic scale, urban microbiomes can exhibit distinct city-specific taxonomic signatures while retaining a shared set of widespread microbial taxa (Danko et al., 2021). Such shared assemblages can be examined through the concept of a core microbiome, defined by the consistent occurrence of microbial taxa across samples or environments (Shade & Handelsman, 2012; Neu et al., 2021). However, taxonomic turnover does not necessarily correspond to equivalent phylogenetic differentiation, as microbial traits can exhibit varying degrees of phylogenetic conservation (Martiny et al., 2013, 2015). Together, these studies highlight the importance of examining microbial communities across spatial scales and raise questions about whether geographic taxonomic turnover in SABs is accompanied by broader phylogenetic differentiation or the persistence of shared bacterial assemblages.

Historic marble monuments provide exposed habitats for SABs in urban environments, where differences in surface texture and in the frequency and duration of wetting may influence community composition. The objectives of this study were to (1) characterize SAB community composition across geographically separated historic marble monuments, (2) determine whether taxonomic turnover is accompanied by broader phylogenetic and inferred ecological differentiation, (3) identify shared and monument-specific core assemblages across monuments, (4) quantify spatial organization at regional (among cities) and local (among sampling locations within monuments) levels, and (5) assess whether core OTUs identified at the two primary monuments were also detected at an independently sampled monument. We hypothesized that SAB communities would exhibit substantial taxonomic turnover among geographically separated monuments while retaining broader phylogenetic and inferred ecological similarities, consistent with the shared constraints of exposed stone environments.

## Materials and methods

### Sample collection

Samples were collected from the rooftops of the Federal Hall National Memorial (New York, NY; hereafter FHNY; March 18, 2024), the Lincoln Memorial (Washington, DC; hereafter LMDC; October 30-31, 2024), and the Merchants’ Exchange Building (Philadelphia, PA, hereafter MEPA; January 29, 2025; Figure 1). These study sites represent three urban marble buildings distributed across the northeastern United States, with New York and Washington, DC serving as the primary comparison sites and Philadelphia providing an intermediate location for independent evaluation. Biofilm material was scraped from the stone surface using a sterile razor blade and transferred into 1.5 mL microcentrifuge tubes containing RLT Plus buffer (Qiagen, Germany) supplemented with β-mercaptoethanol. Samples were immediately placed on dry ice, transported to the laboratory, and stored at −80 °C until nucleic acid extraction.

**Figure 1.**
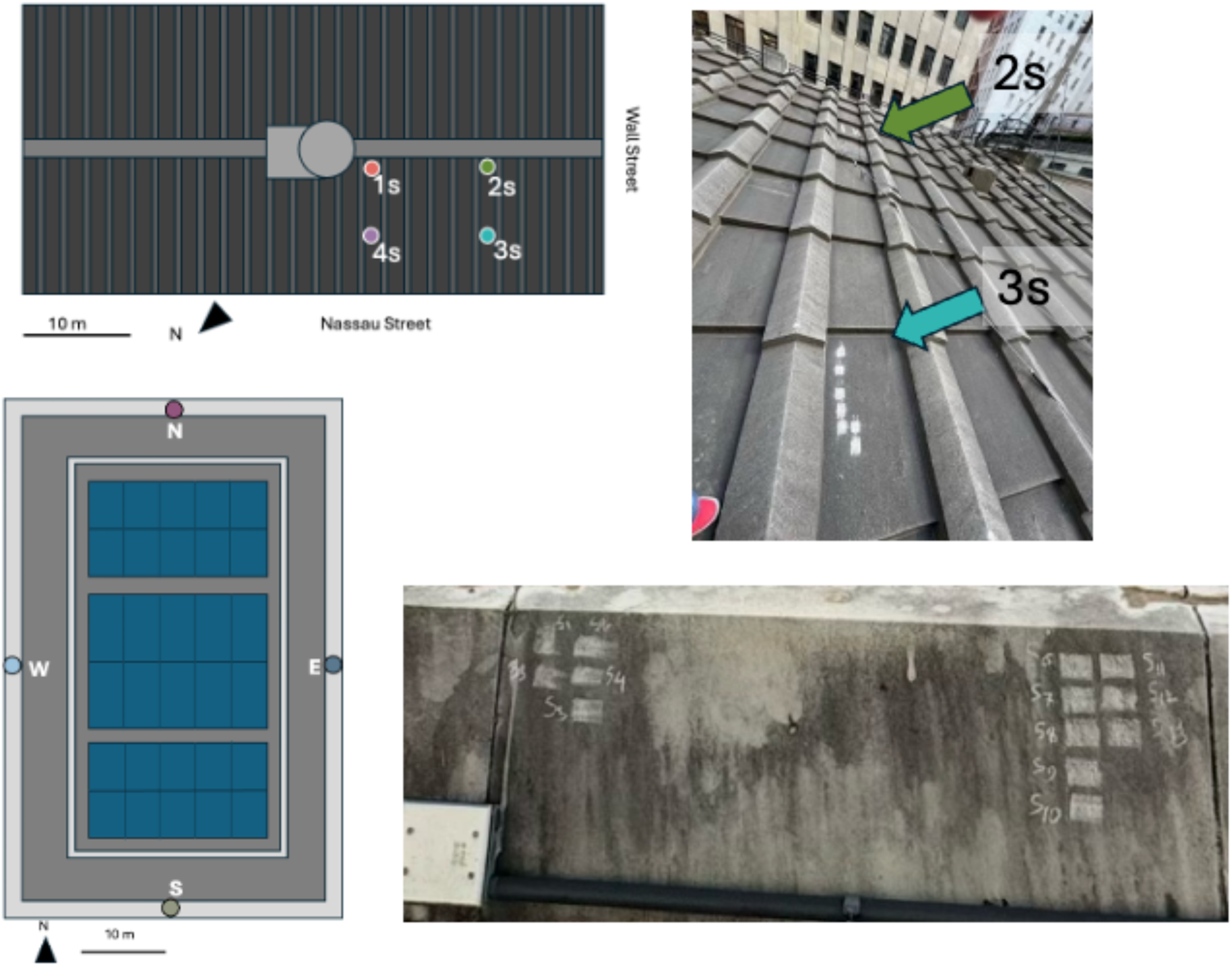
Sampling design at the two marble monuments. Left panels show simplified schematics of the Federal Hall National Memorial (FHNY) roof in New York City and the Lincoln Memorial (LMDC) roof in Washington, DC. Colored dots indicate the four sampling locations (same as Figure 5). Scale bars indicate 10 m. Right panels show representative photographs of the sampling surfaces at each monument. The upper photograph shows scraping locations 2s (top) and 3s (bottom) at FHNY. The lower photograph shows the south sampling location (S) at LMDC.

### Nucleic acid extraction and amplicon sequencing

RNA was extracted and purified using the AllPrep DNA/RNA kit (Qiagen, Germany). Zircon beads were added during extraction to ensure efficient bacterial cell lysis. RNA was used rather than DNA to minimize contributions from dead cells and extracellular DNA. Extracted RNA was reverse transcribed into cDNA using SuperScript III reverse transcriptase and random hexamers (Invitrogen, CA, USA). The V4-V5 region of the bacterial small-subunit rRNA gene was amplified from cDNA using Q5 High-Fidelity DNA Polymerase (New England Biolabs, MA, USA) with the primer pair 515Y (5′-GTG YCA GCM GCC GCG GTA A-3′) and 926R (5′-CCG YCA ATT YMT TTR AGT TT-3′; D’Amore et al., 2016), modified with Illumina adapter sequences (Klindworth et al., 2013). Each sample was amplified in triplicate PCR reactions, which were pooled prior to library preparation to reduce amplification bias (Lahr & Katz, 2009). PCR clean-up, amplicon quantification, barcode addition, library pooling, and Illumina sequencing were performed by the RI-INBRE Molecular Informatics Core.

### Amplicon sequence processing

Illumina MiSeq paired-end reads were processed using a custom Python pipeline developed for both prokaryotic and eukaryotic small-subunit (SSU) rRNA amplicon datasets as described in Sisson et al. (2018) and Grattepanche et al. (2019, 2022) and available at https://github.com/jeandavidgrattepanche/. The workflow performs quality filtering, paired-end merging, primer trimming, dereplication, chimera removal, OTU clustering with SWARM3 (Mahé et al., 2022), OTU table construction, and taxonomic assignment. Intermediate quality-control statistics are generated throughout the pipeline to monitor read retention and clustering performance. Paired-end reads were merged using BBMerge Auto in loose mode with error correction and iterative read extension (ecct, extend2=150, iterations=5). Terminal Ns were trimmed, and sequences with remaining Ns or shorter than 400 bp were excluded. Pooled sequences were clustered with Swarm at a clustering distance of 1. OTUs with at least 100 pooled reads were screened for chimeras.

### Core community identification

For the regional core analysis, we retained samples with at least 5,000 reads and used stratified random sampling by sampling location to select 31 samples per monument. We rarefied the selected samples to 5,000 reads per sample in 20 independent replicates. Within each replicate, an OTU was considered present if it had at least one read and accounted for at least 0.05% of reads in that sample. At the selected rarefaction depth, this corresponded to at least three reads. For each monument, we calculated the fraction of samples in which each OTU was present and averaged that fraction across replicates. An OTU qualified as core at a monument if its mean occupancy was at least 28/31 (90.3% occupancy). OTUs that met this criterion at both FHNY and LMDC formed the regional core. OTUs that met the core criterion at only one of the two monuments were classified as monument-specific core OTUs. We assessed their occurrence separately in the independent MEPA samples. Figure S1 shows how core OTU counts and the fraction of reads they represented varied across the settings tested.

### Sensitivity analysis

To evaluate the robustness of the identified core community, we systematically varied the major parameters of the pipeline, including rarefaction depth, number of rarefaction replicates, occupancy threshold (or maximum allowed absences), minimum number of samples per group, minimum read count, and minimum relative abundance. For each parameter set, the complete workflow was rerun, and the numbers of global and group-specific core OTUs were recorded. In addition to core richness, we quantified the proportion of rarefied sequencing reads represented by each core category, allowing assessment of whether parameter changes primarily affected rare taxa or also altered the dominant community.

### Validation of the regional core

The regional core community was identified using the FHNY and LMDC datasets only. OTUs classified as Global Core or site-specific core were subsequently evaluated in the independent MEPA dataset to determine whether core taxa identified from the two terminal sites were also consistently detected at the intermediate location.

### Taxonomic and phylogenetic analyses

Taxonomic assignments were performed on bacterial OTUs using ARB SILVA Database v128 (Quast et al., 2013) and VSEARCH (Rognes et al., 2016). Taxonomy was assigned to eligible OTUs with more than 100 pooled reads using VSEARCH with minimum thresholds of 97% sequence identity and 80% query coverage. To investigate phylogenetic relationships among core OTUs, representative sequences were combined with reference sequences from ARB SILVA and aligned using MAFFT with default parameters (Katoh & Standley, 2013). Phylogenetic trees were reconstructed using FastTree under the GTR + I model (Price et al., 2010). Core OTUs were categorized according to their taxonomic affiliations and broadly inferred ecological characteristics (Table 2). These ecological categories were intended to provide general ecological context rather than direct functional annotation, as ecological functions were not experimentally verified.

### Statistics

Taxonomic and phylogenetic patterns of bacterial community composition were investigated using nonmetric multidimensional scaling (NMDS) based on Bray–Curtis and weighted UniFrac dissimilarities, respectively (Bray & Curtis, 1957; Lozupone et al., 2007). Bray–Curtis dissimilarities were calculated from OTU abundances, treating individual OTUs as independent taxonomic units, whereas weighted UniFrac incorporated phylogenetic relationships among OTUs and their relative abundances. Differences in community composition among cities and sampling locations were evaluated using permutational multivariate analysis of variance (PERMANOVA; Anderson, 2001). Differences in multivariate dispersion were assessed using PERMDISP (Anderson, 2006) to evaluate whether significant PERMANOVA results could be influenced by differences in within-group variability. Analyses were conducted using the vegan package (Oksanen et al., 2026) in R (R Core Team, 2025), with statistical significance evaluated using 999 permutations.

To characterize environmental variation among sampling locations, principal component analysis (PCA) was performed on standardized environmental measurements, including sampling time, air temperature, stone surface temperature, and relative humidity. Sampling time was expressed as decimal hours. At Federal Hall National Memorial (FHNY), associations between bacterial community composition and environmental variables were further investigated using distance-based redundancy analysis (dbRDA; Legendre & Anderson, 1999; McArdle & Anderson, 2001) based on Bray–Curtis dissimilarities. An initial dbRDA model evaluated the combined association of sampling time, air temperature, stone surface temperature, and relative humidity with community composition. A partial dbRDA was subsequently performed to evaluate the effect of sampling location after conditioning on these environmental variables. Statistical significance was assessed using permutation tests (999 permutations).

## Results

### 1. Dataset overview and bacterial community composition

After quality filtering and removal of low-depth samples (<5,000 reads), 80 samples were retained for downstream analyses. From this dataset, rarefaction to 6,419 reads per sample retained 48.97% of the observed OTUs while preserving 29.72% of the total sequencing reads (Table 1). Rarefaction curves approached saturation for most samples.

**Table 1:**
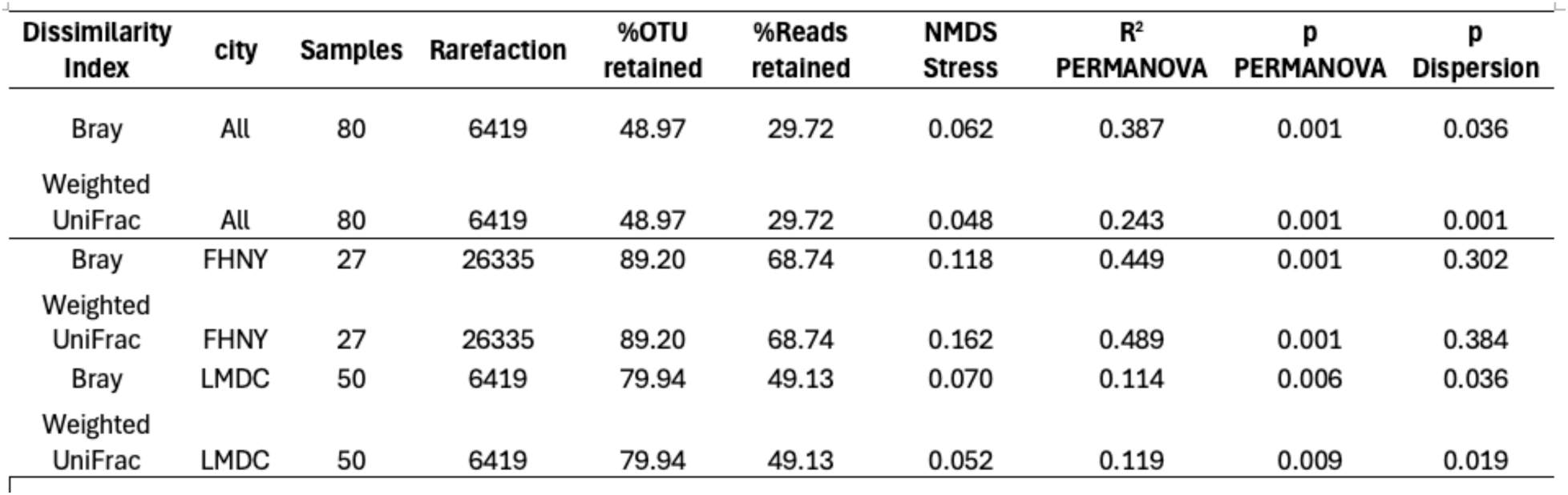
Summary of sequencing, rarefaction, and beta-diversity analyses. For each dataset, the number of samples, rarefaction depth, OTU and read retention, NMDS stress, PERMANOVA (*R*² and *p*), and PERMDISP (*p*) are shown for Bray–Curtis and weighted UniFrac dissimilarities.

| **Dissimilarity Index** | **city** | **Samples** | **Rarefaction** | **%OTU retained** | **%Reads retained** | **NMDS Stress** | **R<sup>2</sup> PERMANOVA** | **p PERMANOVA** | **p Dispersion** |
|---|---|---|---|---|---|---|---|---|---|
| Bray | All | 80 | 6419 | 48.97 | 29.72 | 0.062 | 0.387 | 0.001 | 0.036 |
| Weighted UniFrac | All | 80 | 6419 | 48.97 | 29.72 | 0.048 | 0.243 | 0.001 | 0.001 |
| Bray | FHNY | 27 | 26335 | 89.20 | 68.74 | 0.118 | 0.449 | 0.001 | 0.302 |
| Weighted UniFrac | FHNY | 27 | 26335 | 89.20 | 68.74 | 0.162 | 0.489 | 0.001 | 0.384 |
| Bray | LMDC | 50 | 6419 | 79.94 | 49.13 | 0.070 | 0.114 | 0.006 | 0.036 |
| Weighted UniFrac | LMDC | 50 | 6419 | 79.94 | 49.13 | 0.052 | 0.119 | 0.009 | 0.019 |

The 100 most abundant OTUs represented the majority of reads in each sample and revealed clear differences in community composition among the three monuments (Figure 2A). The bacterial communities were dominated by members of the Proteobacteria, Cyanobacteria, Actinobacteriota, Bacteroidota, Chloroflexi, and Acidobacteriota, although their relative abundances varied substantially among samples (Order-level profiles are shown in Figure 2B). Samples from FHNY were characterized by relatively consistent taxonomic profiles, whereas LMDC exhibited greater variability among individual samples. The three MEPA samples displayed intermediate community composition, sharing abundant taxa with both FHNY and LMDC (Figure 2).

**Figure 2.**
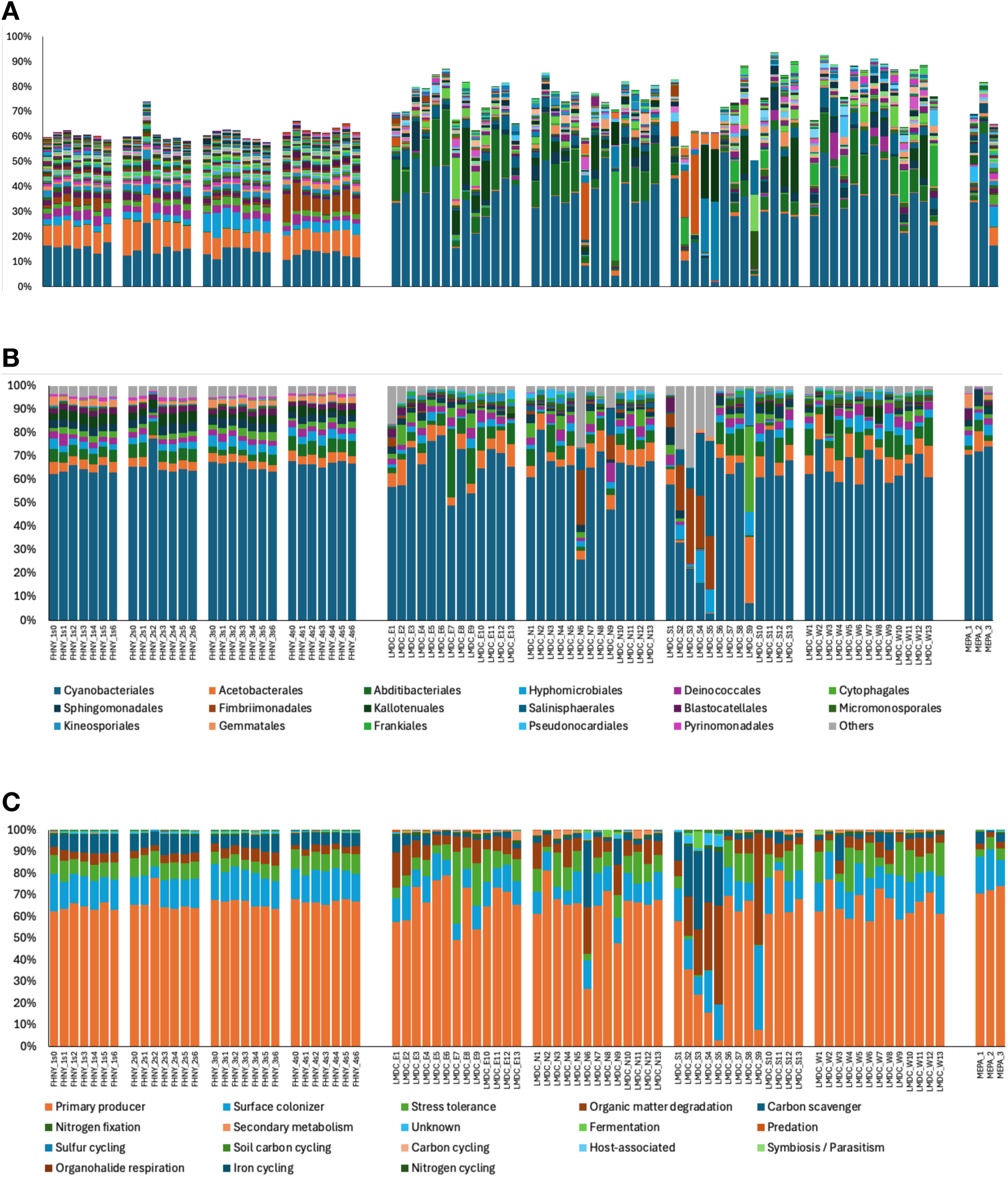
Community (A and B) composition and (C) function. For the composition, each color represents one OTU among the 100 most abundant OTUs (A) or an order (B). Orders with a mean relative abundance below 0.5% were grouped as “Others”. (C) Functions are inferred based on the order of each OTU.

### 2. Large-scale biogeography

Patterns of bacterial community composition were first examined among the three marble monuments using Bray–Curtis and weighted UniFrac dissimilarity metrics (Figure 3; Table 1). Bray–Curtis NMDS revealed clear taxonomic differentiation among cities (PERMANOVA *R*² = 0.387, *P* = 0.001), with FHNY and LMDC forming distinct clusters and the Philadelphia samples (MEPA) occupying an intermediate position between the two cities. Multivariate dispersion differed significantly among cities (PERMDISP *P* = 0.036), reflecting greater variability within LMDC than within FHNY (Figure 3).

**Figure 3.**
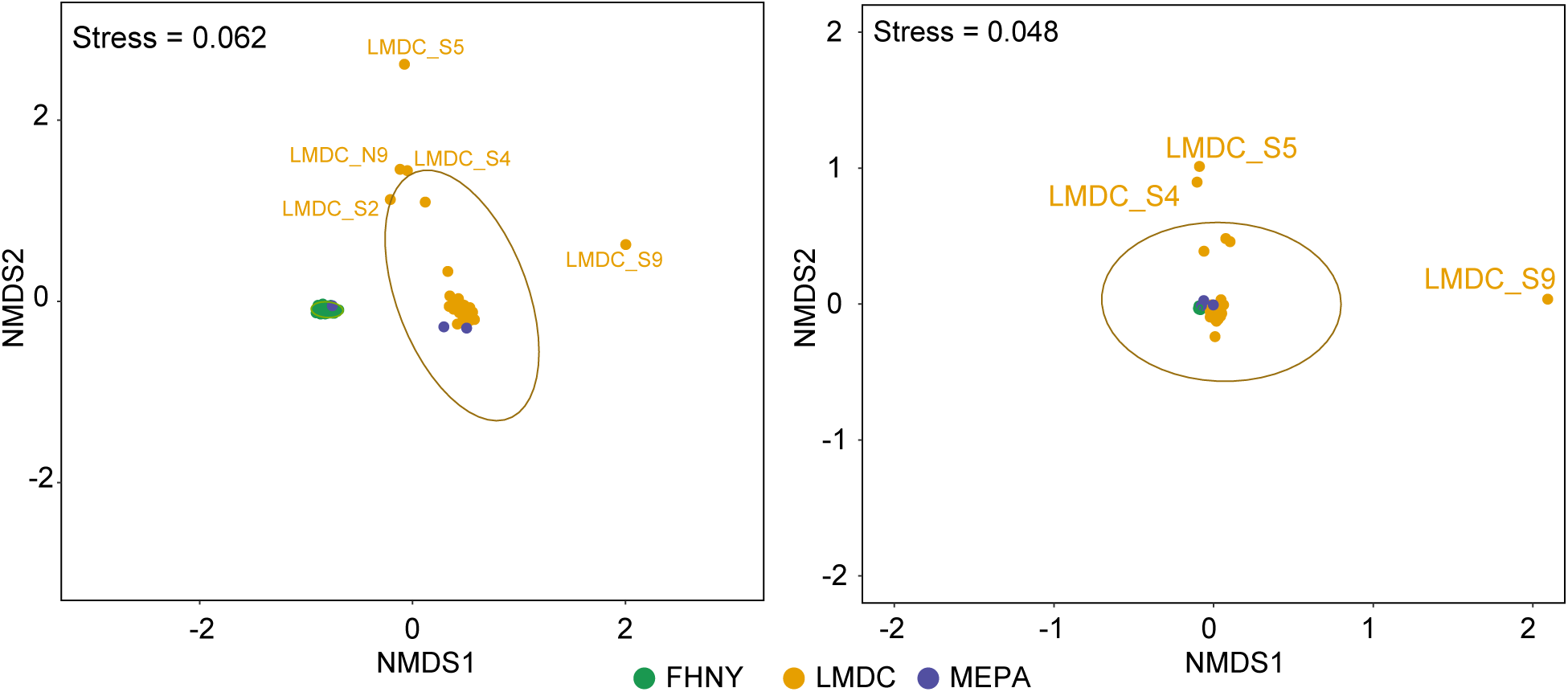
Community structure inferred by non-metric multidimensional scaling (NMDS). NMDS ordinations based on Bray–Curtis dissimilarity (left; stress = 0.062) and weighted UniFrac distance (right; stress = 0.048) for bacterial communities collected from the three marble monuments: Federal Hall National Memorial, New York (FHNY; green), Lincoln Memorial, Washington, DC (LMDC; orange), and Merchants’ Exchange Building, Philadelphia (MEPA; blue). Samples were rarefied to 6,419 reads prior to distance calculation. Ellipses represent the 95% confidence interval for each site. Bray–Curtis dissimilarity reveals greater taxonomic differentiation among sites, whereas weighted UniFrac indicates a higher degree of phylogenetic similarity, particularly between the FHNY and LMDC communities.

In contrast, weighted UniFrac NMDS showed substantially greater overlap among the three cities despite remaining significantly different (PERMANOVA *R*² = 0.243, *P* = 0.001). Most FHNY and MEPA samples clustered closely together, whereas only a few LMDC samples (S4, S5, and S9) remained clearly separated from the main group. Multivariate dispersion also differed among monuments for weighted UniFrac (PERMDISP *P* = 0.001). The lower PERMANOVA *R*² and greater overlap among monuments with weighted UniFrac than with Bray–Curtis were consistent with some taxonomic turnover occurring among closely related bacterial lineages (Figure 3).

Despite the pronounced taxonomic turnover observed among monuments, weighted UniFrac revealed substantially greater overlap among bacterial communities than Bray–Curtis dissimilarity (Figure 3). This pattern indicates that differences in community composition primarily reflected replacement among closely related bacterial lineages rather than turnover among distantly related clades. The inferred ecological profiles were broadly similar across the three monuments. Chlorophyll-containing bacteria accounted for the largest fraction of reads, followed by heterotrophic taxa, whereas other ecological strategies collectively accounted for only a minor fraction of the communities (Figure 2C).

### 3. A conserved urban core assemblage

To identify bacterial lineages consistently associated with historic marble monuments, we defined core assemblages based on their prevalence across samples and examined their phylogenetic relationships (Figure 4). Allowing more absences increased the number of OTUs classified as core (Figure S1), and we used a three-absence allowance for the regional analysis. Although individual OTUs differed markedly among monuments, numerous bacterial lineages were shared across geographically separated sites. The core assemblage comprised 92 OTUs, including 15 OTUs shared among the two monuments, 64 OTUs specific to FHNY (hereafter FHNY-specific core), and 13 OTUs specific to LMDC (hereafter LMDC-specific core; Figure 4).

**Figure 4.**
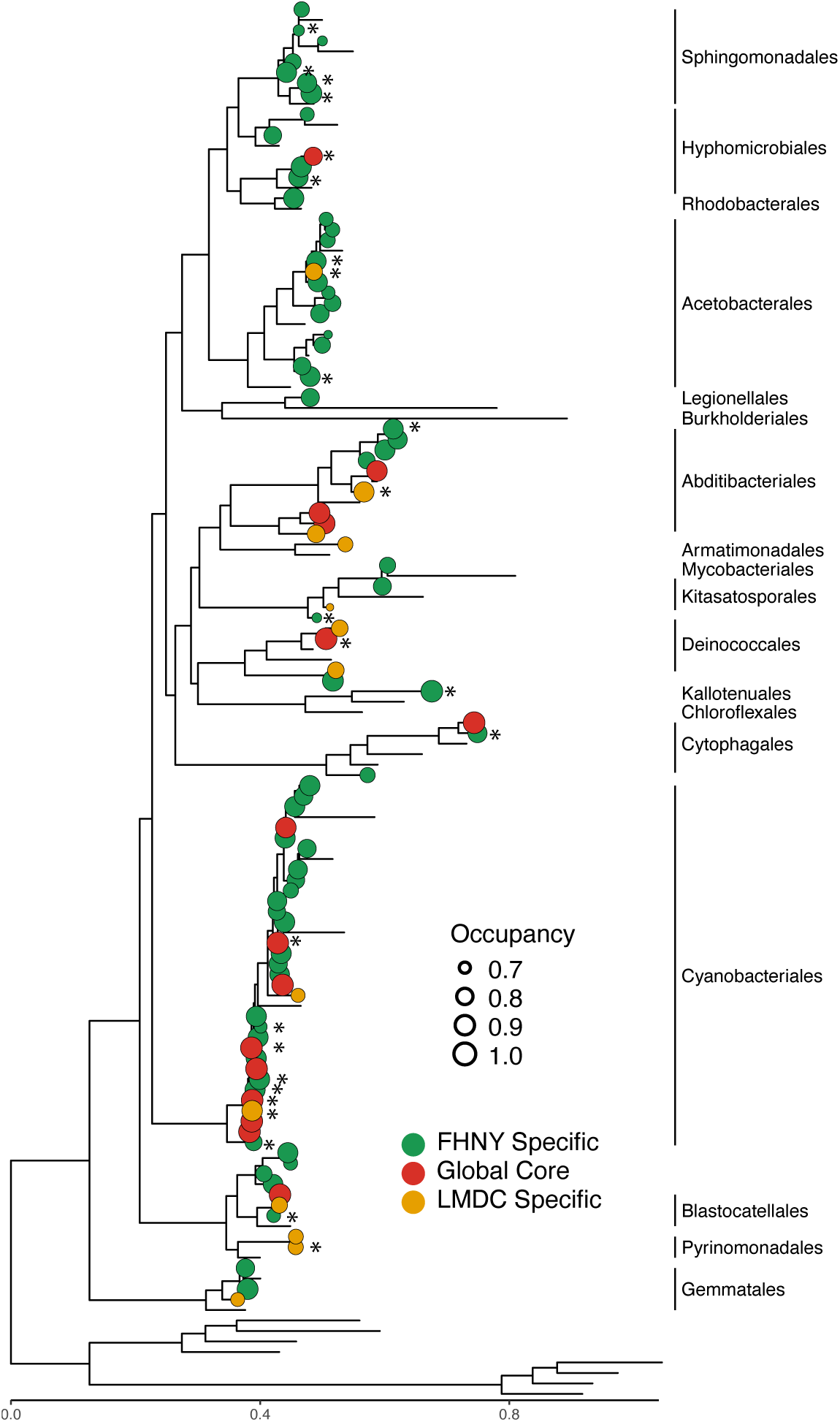
Phylogenetic relationships among global and site-specific core bacterial lineages. An asterisk (*) denotes core lineages that were additionally present in all three Merchants’ Exchange Building (MEPA) samples.

The global core assemblage comprised representatives from several phylogenetically diverse bacterial groups, including Cyanobacteriales, Hyphomicrobiales, Deinococcales, and multiple Acidobacteriota lineages (Figure 4). These taxa encompassed a broad range of ecological strategies, including oxygenic phototrophs, oligotrophic surface colonizers, stress-tolerant heterotrophs, and biofilm-forming bacteria (Table 2).

**Table 2:** Reported traits and typical metabolisms of representative taxa within bacterial orders detected in the core assemblages to provide ecological context for the conserved bacterial lineages identified in Figure 4. The relevance column describes how these traits could contribute to occurrence in marble biofilms or other built environments. These roles were inferred from the literature.

| **Order** | **Reported trait** | **Typical metabolism** | **Potential relevance to marble / built environments** | **References** |
|---|---|---|---|---|
| Hyphomicrobiales | Methylotrophy in some members | Heterotrophy | Use of one-carbon compounds when available | Hördt et al. (2020); Urakami et al. (1995) |
| Rhodobacterales | Light harvesting in some roseobacters | Heterotrophy or photoheterotrophy | Supplemental light energy on exposed surfaces | Luo & Moran (2014) |
| Sphingomonadales | Aromatic compound degradation | Aerobic heterotrophy | Use of deposited aromatic compounds | White et al. (1996); Stolz (2009) |
| Acetobacterales | Acetic acid production in some members | Aerobic oxidation of alcohols and sugars | Local acidification where substrates are available | Hördt et al. (2020); Guzman & Vilcinskas (2022) |
| Burkholderiales | Diverse substrate use in Burkholderiaceae | Heterotrophy | Use of varied organic inputs | Coenye (2014) |
| Legionellales | Intracellular growth in protozoa | Host-associated heterotrophy | Persistence in biofilm-associated protists | Newton et al. (2010) |
| Abditibacteriales | Growth on dilute media in a cultured representative | Aerobic heterotrophy | Growth where organic nutrients are dilute | Tahon et al. (2018) |
| Armatimonadales | Use of plant-derived polysaccharides | Aerobic heterotrophy | Use of plant-derived material deposited on surfaces | Tamaki et al. (2011) |
| Kitasatosporales | Secondary metabolite production | Heterotrophy | Potential chemical interactions with neighboring microbes | Nouioui et al. (2018); Takahashi (2017) |
| Mycobacteriales | Biofilm association in some mycobacteria | Aerobic heterotrophy | Persistence on intermittently wet built surfaces | Falkinham et al. (2001); Tortoli et al. (2017) |
| Deinococcales | UV and desiccation tolerance | Aerobic heterotrophy | Survival during drying and UV exposure | Slade & Radman (2011) |
| Chloroflexales | Light harvesting in some members | Heterotrophy, with anoxygenic phototrophy in some members | Supplemental light energy in illuminated biofilms | Ward et al. (2018) |
| Kallotenuales | Cellulose degradation in the type strain | Aerobic heterotrophy | Potential use of cellulose-rich debris | Cole et al. (2013); Parks et al. (2018) |
| Cytophagales | Polymer degradation in some Cytophagaceae | Predominantly aerobic heterotrophy | Breakdown of organic polymers in the biofilm | McBride et al. (2014) |
| Cyanobacteriales | Photosynthetic carbon fixation | Oxygenic photosynthesis | Organic carbon production on illuminated marble | Whitton & Potts (2000) |
| Blastocatellales | Cultured representatives from semiarid soil | Aerobic heterotrophy | Potential persistence on periodically dry surfaces | Foesel et al. (2013) |
| Pyrinomonadales | Growth on organic substrates in a cultured representative | Aerobic heterotrophy | Use of organic carbon produced within the biofilm | Crowe et al. (2014) |
| Gemmatales | Carbohydrate use in *Gemmata* | Aerobic heterotrophy | Use of biofilm-derived carbohydrates | Fuerst & Sagulenko (2011); Fuerst et al. (2022) |

Comparison of city-specific core assemblages showed substantial phylogenetic overlap despite differences in OTU composition. Several major bacterial orders were shared among all monuments, whereas a smaller number of lineages appeared restricted to individual cities. Several lineages were well represented in the FHNY-specific core assemblage, including Sphingomonadales, some members of which can degrade aromatic compounds (Table 2).

To assess whether the conserved core extended beyond the two extensively sampled monuments, we examined three independent samples collected from the Merchants’ Exchange Building (MEPA) in Philadelphia. Five of the 15 global-core OTUs were also detected in all three MEPA samples (1/3 of the global core assemblage), together with 16 FHNY-core OTUs (25% of the FHNY core) and four LMDC-core OTUs (31% of the LMDC core; Figure 4). Of the five global-core OTUs, three were assigned to Cyanobacteriales, one to Deinococcales, and one to Hyphomicrobiales (Figure 4). Although based on limited sampling, these observations provide independent support for the widespread occurrence of several conserved bacterial lineages across historic urban marble monuments. Collectively, these results indicate that geographically separated marble monuments support a conserved phylogenetic core despite substantial taxonomic turnover at finer taxonomic resolution.

### 4. Contrasting biogeography within marble monuments

To evaluate bacterial community biogeography at the “within-monument” scale, ordinations were repeated separately for FHNY and LMDC using sampling locations as the grouping factor (Figure 5; Table 1). Bray–Curtis and weighted UniFrac dissimilarities were used to distinguish taxonomic from phylogenetic patterns, as performed previously (Figure 3).

**Figure 5.**
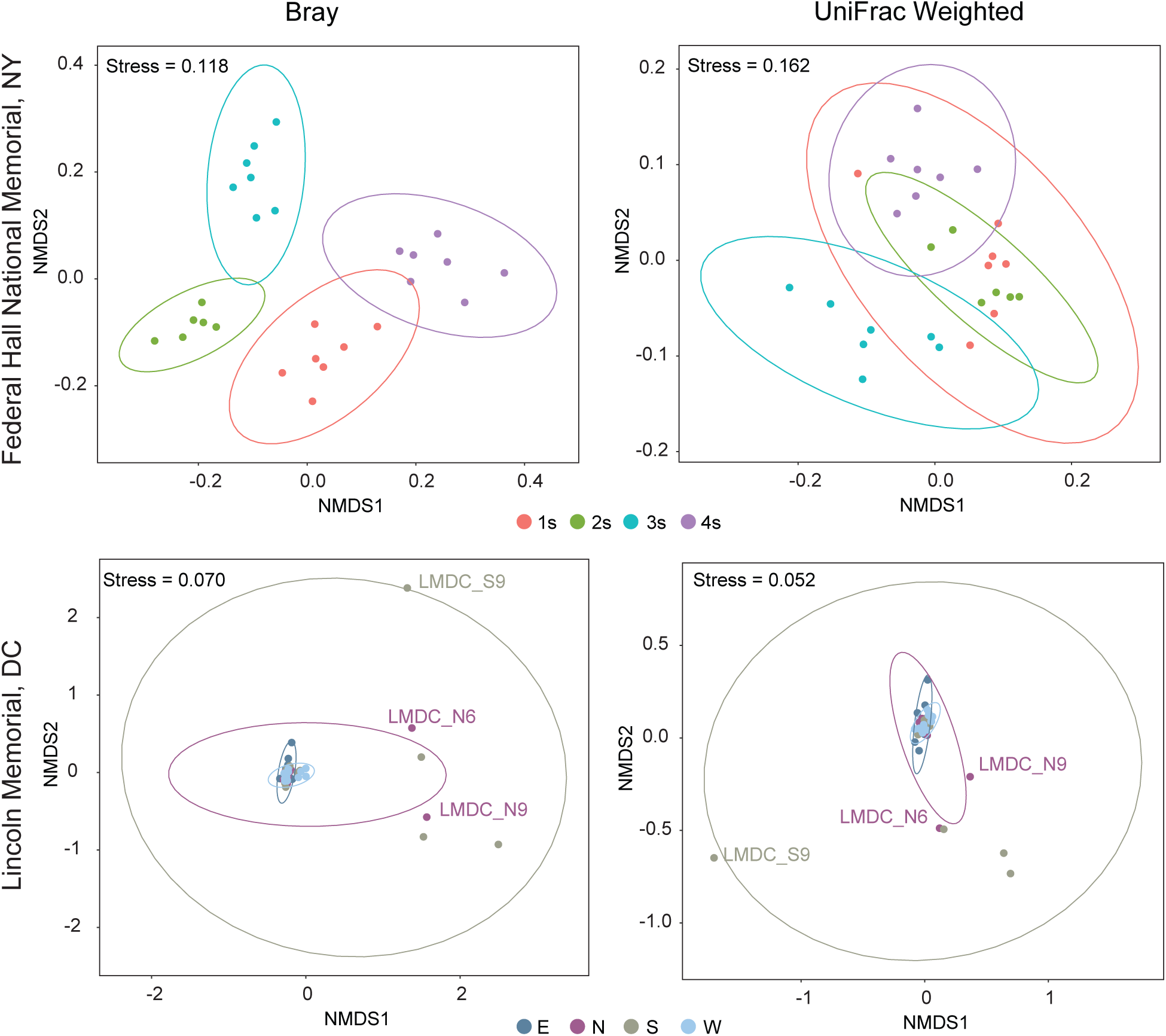
Community structure inferred by non-metric multidimensional scaling (NMDS). NMDS ordinations based on Bray–Curtis dissimilarity (left) and weighted UniFrac distance (right) for bacterial communities collected at Federal Hall National Memorial, NY (Top) and Lincoln Memorial, DC (Bottom). Ellipses represent the 95% confidence interval for each site. Colors match the sampling locations shown in Figure 1.

At FHNY, bacterial communities were strongly structured according to the four sampling locations despite their close spatial proximity on a single roof (Figure 1). Bray–Curtis NMDS resolved four distinct clusters corresponding to the four sampling sites (PERMANOVA *R*² = 0.449, *P* = 0.001), while weighted UniFrac NMDS showed substantial overlap among the same samples despite a significant PERMANOVA (*R*² = 0.489, *P* = 0.001). No significant differences in multivariate dispersion were detected for either Bray–Curtis or weighted UniFrac distances (PERMDISP *P* = 0.302 and 0.384, respectively), indicating that differences among sampling locations primarily reflected shifts in community composition rather than differences in within-site variability.

In contrast, bacterial communities from the four roof orientations at LMDC overlapped extensively in both NMDS ordinations (Figure 5). PERMANOVA detected modest but significant differences among orientations (Bray–Curtis *R*² = 0.114, *P* = 0.006; weighted UniFrac *R*² = 0.119, *P* = 0.009). Significant multivariate dispersion (Bray–Curtis *P* = 0.036; weighted UniFrac *P* = 0.019) reflected increased variability among samples within sampling locations, with a small number of samples contributing disproportionately to the observed heterogeneity (Figure 5).

Together, these results reveal contrasting patterns of small-scale biogeography between the two monuments. At FHNY, samples from locations only a few meters apart formed distinct clusters in the Bray–Curtis ordination, whereas at LMDC, samples from the four roof orientations overlapped extensively and showed significant differences in dispersion among orientations. Environmental measurements tended to group by monument in the PCA, while the FHNY-only PCA showed no clear grouping by sampling location (Figure S2). The combined environmental variables (sampling time, air temperature, stone surface temperature, and relative humidity) showed a marginal but nonsignificant association with community composition at FHNY (dbRDA, R² = 0.194, adjusted R² = 0.048, P = 0.062). Importantly, spatial differentiation remained significant after accounting for these variables (partial dbRDA, F = 5.633, P = 0.001; Figure S3).

## Discussion

Subaerial bacterial communities colonizing historic marble monuments exhibited pronounced taxonomic turnover across geographically separated sites while maintaining a remarkably conserved phylogenetic and inferred ecological composition. Despite differences in OTU composition among monuments, bacterial communities were consistently dominated by closely related lineages and shared a conserved core assemblage comprising phototrophic, oligotrophic, and stress-tolerant taxa. Fine-scale community organization also differed markedly between monuments, indicating that the ecological processes shaping bacterial communities vary across spatial scales. Together, these observations provide insight into how environmental filtering and local ecological processes contribute to the biogeography of bacterial communities on historic urban marble surfaces.

### Conserved core assemblages

The conserved core assemblage suggests that geographically separated marble monuments select for bacterial lineages sharing ecological traits suited to exposed stone surfaces. Despite differences in taxonomic composition, the core consistently included phototrophic Cyanobacteriales, oligotrophic Alphaproteobacteria (e.g., Hyphomicrobiales and Sphingomonadales), stress-tolerant Deinococcales, and several Acidobacteriota lineages (Table 2). Similar bacterial groups have been reported from subaerial biofilms colonizing sandstone and limestone sculptures (Cattò et al., 2023), as well as natural rock outcrops, desert crusts, and other exposed lithic environments, where microorganisms experience chronic nutrient limitation, intense solar radiation, and repeated desiccation (Gaylarde, 2020; Macedo et al., 2009; Villa et al., 2016; Villa & Cappitelli, 2019). Together, these groups encompass complementary ecological strategies that likely facilitate persistence on nutrient-poor, highly irradiated, and periodically desiccated mineral surfaces (see Table 2 for ecological strategies and references). Rather than representing monument-specific taxa, the conserved core appears to reflect a common set of bacterial lineages adapted to the environmental constraints characteristic of urban marble habitats.

On exposed marble surfaces, phototrophic Cyanobacteriales likely establish primary production, while heterotrophic bacteria exploit locally available organic substrates and contribute to mature biofilm development (Table 2). A recent model proposes that phototrophs release excess fixed carbon, providing a substrate that can support heterotrophic growth on exposed marble (Tenore et al., 2026). Different OTUs may fill similar ecological roles at different monuments, allowing complementary processes such as primary production and the use of organic carbon to recur despite taxonomic turnover. Among these heterotrophs, some members of the Sphingomonadales can degrade aromatic compounds, a trait that may allow them to use organic material deposited on exposed marble surfaces (Table 2). UV and desiccation tolerance in some Deinococcales may help them persist on exposed marble, while members of Gemmatales and some Acidobacteriota lineages may use organic substrates available within the biofilm (Table 2). Together, these complementary ecological strategies likely facilitate the persistence of a conserved bacterial core on nutrient-poor marble surfaces despite substantial taxonomic turnover.

The independent detection of many core lineages in the Merchants’ Exchange Building (MEPA) further suggests that these taxa are not restricted to a single monument but may represent widespread colonizers of urban marble monuments (Figure 4). Although additional sites and longitudinal sampling will be required to determine the geographic extent and temporal stability of this core, the recurrence of similar bacterial lineages across geographically separated marble monuments is consistent with the influence of shared environmental constraints.

### Taxonomic turnover with phylogenetic conservation

Our results show that geographically separated monuments harbor distinct bacterial assemblages at the OTU level, while remaining considerably similar when phylogenetic relationships are taken into account (Figures 2, 3 and 5). The most striking finding of this study was the contrast between strong taxonomic turnover and comparatively limited phylogenetic differentiation among monuments. Communities from different monuments overlapped more when we accounted for evolutionary relationships among bacteria (weighted UniFrac) than when we compared OTU abundances alone (Bray–Curtis, Figure 3). This pattern suggests that some of the OTUs that differed among monuments belonged to closely related bacterial lineages. The monuments also shared a set of core OTUs (Figure 4) and had broadly similar inferred ecological profiles (Figure 2C). The recurrence of related lineages and similar inferred ecological profiles across monuments suggests that the conditions on exposed marble may favor bacteria with similar ecological traits, even as individual OTUs vary among monuments.

Similar patterns of environmental filtering have been reported across a wide range of microbial and macrobial systems, where environmental conditions select for organisms sharing ecological traits rather than identical taxa (Webb et al., 2002; Cavender-Bares et al., 2009). Because some microbial traits are shared among closely related lineages, similar environmental conditions may favor related bacteria even when the particular OTUs differ among monuments (Martiny et al., 2013, 2015).

Subaerial biofilms develop under persistent environmental constraints, including chronic nutrient limitation, repeated desiccation, intense solar radiation, and large temperature fluctuations, all of which restrict the range of bacterial lineages capable of successfully colonizing and persisting on exposed stone (Villa et al., 2016; Villa & Cappitelli, 2019). Rather than selecting for individual taxa, these environmental conditions likely favor broader ecological characteristics, such as phototrophy, oligotrophy, biofilm formation, and tolerance to desiccation and UV exposure. Consequently, different, but closely related, bacterial lineages possessing similar ecological strategies may establish at different monuments, resulting in substantial OTU turnover while maintaining a comparatively conserved phylogenetic framework.

This interpretation is consistent with the widespread occurrence of closely related bacterial groups across natural-rock outcrops, desert crusts, and stone monuments, where exposure to similar environmental pressures repeatedly selects for microorganisms adapted to life on mineral surfaces (Villa & Cappitelli, 2019). Although ecological traits were inferred from taxonomic identity rather than direct measurements, the similarities in phylogenetic organization and inferred ecological compositions are consistent with the idea that geographically separated marble monuments impose similar environmental constraints on their bacterial communities.

### Local ecological heterogeneity

While environmental filtering appears to constrain the broader phylogenetic organization of bacterial communities across monuments, analyses within individual monuments revealed that local ecological processes can generate markedly different patterns of spatial organization. Contrary to expectation, bacterial communities at FHNY exhibited stronger fine-scale spatial organization than those at LMDC, despite the four sampling sites being separated by only a few meters on a single roof surface. This finding indicates that broad architectural characteristics alone cannot explain the spatial organization of subaerial bacterial communities. The contrasting spatial organization observed between monuments suggests that ecological processes operating at the microscale differ substantially among historic marble surfaces. At FHNY, community composition remained associated with sampling location after accounting for sampling time, air temperature, stone surface temperature, and relative humidity. Local differences in shading, reflected light, or runoff along the roof slope could contribute to this pattern. At LMDC, bacterial communities from the four roof orientations overlapped extensively in both ordinations, although the orientations differed in how variable their communities were. Roof orientation therefore provided a limited basis for distinguishing community composition at LMDC. Differences in surface exposure and colonization history could help explain these patterns at both monuments. Some factors impacting SAB growing on other stone materials include surface exposure, position, and geometry (Cattò et al., 2023). Together, these observations suggest that localized ecological conditions may outweigh broader architectural features in structuring bacterial communities on exposed stone surfaces.

Viewed from this perspective, the significant multivariate dispersion observed at LMDC represents more than a statistical property of the data; it reflects genuine ecological heterogeneity among neighboring biofilms. Such heterogeneity could arise from localized nutrient deposition, differential water retention, stochastic colonization, or differences in biofilm succession, all of which operate at spatial scales smaller than those captured by the sampling design. Distinguishing among these processes will require longitudinal sampling coupled with direct environmental measurements, but the contrasting patterns observed at FHNY and LMDC clearly demonstrate that microscale spatial organization is not governed solely by the physical layout of the monument. Together, these observations indicate that bacterial communities inhabiting historic marble monuments are structured by ecological processes operating across multiple spatial scales, from broad environmental filtering among monuments to localized processes generating heterogeneity within individual biofilms.

## Conclusions

Historic urban marble monuments harbor taxonomically distinct bacterial communities while maintaining a conserved phylogenetic framework and a shared core assemblage across geographically separated sites. These findings suggest that bacterial biogeography on exposed stone surfaces is shaped by environmental filtering operating across broad spatial scales together with localized ecological processes acting within individual monuments. Whether these localized assemblages represent persistent spatial niches or transient communities remains unknown. Longitudinal sampling will be required to determine the temporal stability of the conserved phylogenetic framework identified here. Integrating metagenomics, metatranscriptomics, or physiological characterization will further clarify the extent to which the conserved phylogenetic patterns translate into conservation of ecological traits and functional potential.

## Data Accessibility Statement

Raw bacterial SSU rRNA amplicon reads are deposited in the NCBI Sequence Read Archive (SRA) under BioProject PRJNA1560971.

