## Supplementary materials for "Taxonomic turnover with phylogenetic conservation in subaerial biofilm (SAB) communities of marble monuments"

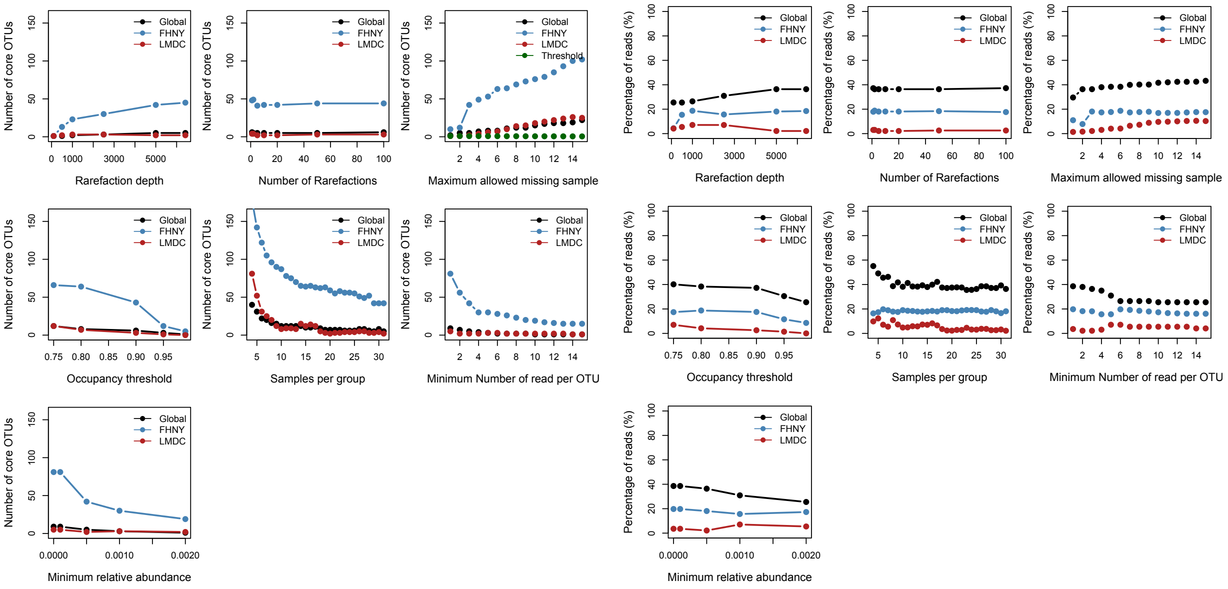


**Figure S1** Sensitivity analysis of core assemblage definition. The number of core OTUs (left panels) and the percentage of sequencing reads represented by the core assemblage (right panels) are shown as a function of the seven parameters used to define the core: rarefaction depth, number of rarefactions, maximum allowed missing samples, minimum number of samples per group, occupancy threshold, minimum number of reads per OTU, and minimum relative abundance. Results are presented for global core assemblage (Global core shared by FHNY and LMDC samples), the Federal Hall National Memorial core assemblage (FHNY; all FHNY samples), and the Lincoln Memorial core assemblage (LMDC; all LMDC samples).

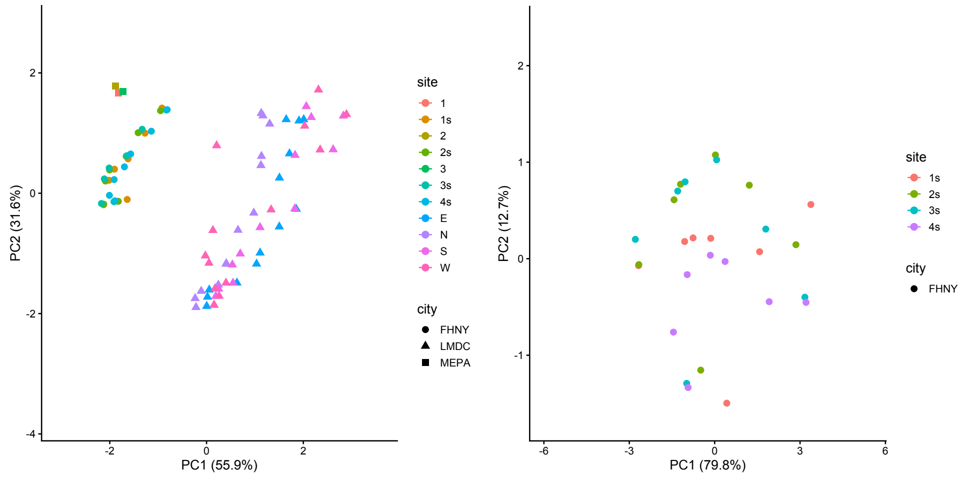


**Figure S2** Environmental variation across sampling locations. Environmental PCA across FHNY, LMDC, and MEPA, colored by sampling location and shaped by city (left); or restricted to FHNY, colored by sampling location (right).

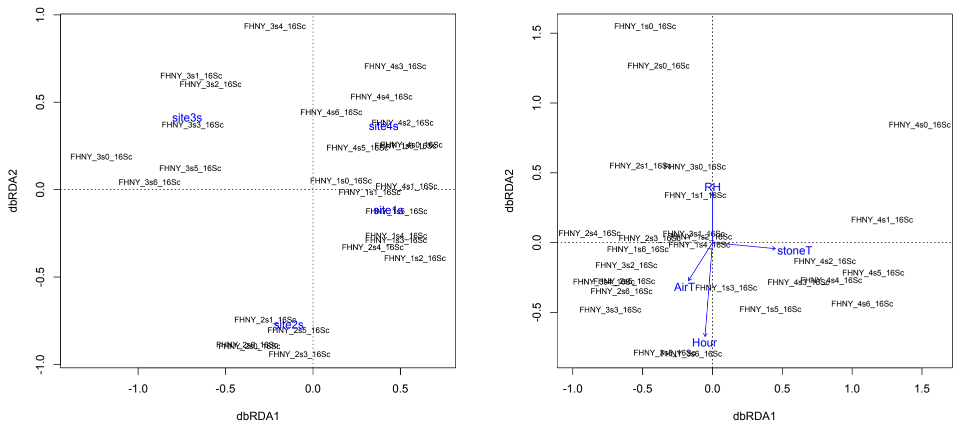


**Figure S3** Distance-based redundancy analyses (dbRDA) of bacterial community composition at Federal Hall National Memorial (FHNY), based on Bray–Curtis dissimilarities. Left - Partial dbRDA showing differences among sampling locations after accounting for four measured environmental parameters (P = 0.001). Right - dbRDA showing associations between bacterial community composition and the measured environmental variables (P = 0.062).
